# Mechanism underlying the DNA-binding preferences of the *Vibrio cholerae* and vibriophage VP882 VqmA quorum-sensing receptors

**DOI:** 10.1101/2021.04.15.439952

**Authors:** Olivia P. Duddy, Xiuliang Huang, Justin E. Silpe, Bonnie L. Bassler

## Abstract

Quorum sensing is a chemical communication process that bacteria use to coordinate group behaviors. In the global pathogen *Vibrio cholerae*, one quorum-sensing receptor and transcription factor, called VqmA (VqmA_Vc_), activates expression of the *vqmR* gene encoding the small regulatory RNA VqmR, which represses genes involved in virulence and biofilm formation. Vibriophage VP882 encodes a VqmA homolog called VqmA_Phage_ that activates transcription of the phage gene *qtip*, and Qtip launches the phage lytic program. Curiously, VqmA_Phage_ can activate *vqmR* expression but VqmA_Vc_ cannot activate expression of *qtip*. Here, we investigate the mechanism underlying this asymmetry. We find that promoter selectivity is driven exclusively by each VqmA DNA-binding domain and key DNA sequences in the *vqmR* and *qtip* promoters are required to maintain specificity. A protein sequence-guided mutagenesis approach revealed that the residue E194 of VqmA_Phage_ and A192, the equivalent residue in VqmA_Vc_, in the helix-turn-helix motifs contribute to promoter-binding specificity. A genetic screen to identify VqmA_Phage_ mutants that are incapable of binding the *qtip* promoter but maintain binding to the *vqmR* promoter delivered additional VqmA_Phage_ residues located immediately C-terminal to the helix-turn-helix motif as required for binding the *qtip* promoter. Surprisingly, these residues are conserved between VqmA_Phage_ and VqmA_Vc_. A second, targeted genetic screen revealed a region located in the VqmA_Vc_ DNA-binding domain as necessary to prevent VqmA_Vc_ from binding the *qtip* promoter, thus restricting DNA-binding to the *vqmR* promoter. We propose that the VqmA_Vc_ helix-turn-helix motif and the C-terminal flanking residues function together to prohibit VqmA_Vc_ from binding the *qtip* promoter.

**AUTHOR SUMMARY:** Bacteria use a chemical communication process called quorum sensing (QS) to orchestrate collective behaviors. Recent studies demonstrate that bacteria-infecting viruses, called phages, also employ chemical communication to regulate collective activities. Phages can encode virus-specific QS-like systems, or they can harbor genes encoding QS components resembling those of bacteria. The latter arrangement suggests the potential for chemical communication across domains, i.e., between bacteria and phages. Ramifications stemming from such cross-domain communication are not understood. Phage VP882 infects the global pathogen *Vibrio cholerae*, and “eavesdrops” on *V. cholerae* QS to optimize the timing of its transition from living as a parasite to killing the host, and moreover, to manipulate *V. cholerae* biology. To accomplish these feats, phage VP882 relies on VqmA_Phage_, the phage-encoded homolog of the *V. cholerae* VqmA_Vc_ QS receptor and transcription factor. VqmA_Vc_, by contrast, is constrained to the control of only *V. cholerae* genes and is incapable of regulating phage biology. Here, we discover the molecular mechanism underpinning the asymmetric transcriptional preferences of the phage-encoded and bacteria-encoded VqmA proteins. We demonstrate how VqmA transcriptional regulation is crucial to the survival and persistence of both the pathogen *V. cholerae*, and the phage that preys on it.

## INTRODUCTION

Quorum sensing (QS) is a cell-cell communication process that allows bacteria to coordinate collective behaviors (1). QS relies on the production, release, and group-wide detection of extracellular signaling molecules called autoinducers (AIs). In the global pathogen *Vibrio cholerae*, the AI, 3,5-dimethly-pyrazin-2-ol (DPO), together with its partner cytoplasmic QS receptor and transcription factor, VqmA (VqmA_Vc_), comprises one of the QS circuits that controls group behaviors (2–4). VqmA_Vc_, following binding to DPO, activates transcription of the *vqmR* gene encoding the small RNA, VqmR, which, in turn, represses the expression of genes required for biofilm formation and virulence factor production (2–4).

Recently, bacteria-specific viruses, called phages, have been shown to engage in density-dependent regulation of their lysis-lysogeny decisions via chemical dialogs (5,6). Germane to our studies are phages that encode proteins resembling bacterial QS components (5,7). Vibriophage VP882 is one such phage: It encodes the QS receptor VqmA (VqmA_Phage_), a homolog of the *V. cholerae* QS receptor VqmA_Vc_ (5). VqmA_Phage_, like VqmA_Vc_, binds host-produced DPO. DPO-bound VqmA_Phage_ activates transcription of the phage gene *qtip*. Qtip is an antirepressor that sequesters the phage VP882 repressor of lysis, leading to derepression of the phage lytic program and killing of the *Vibrio* host at high cell density (5,8). Thus, the DPO AI mediates both bacterial and phage lifestyle decisions. Curiously, VqmA_Phage_ can substitute for VqmA_Vc_ to activate the *V. cholerae vqmR* promoter (P*vqmR*) (5). In contrast, VqmA_Vc_ cannot substitute for VqmA_Phage_ and recognize the phage VP882 *qtip* promoter (P*qtip*). Presumably, the ability of VqmA_Phage_ to bind both P*vqmR* and P*qtip* provides phage VP882 the capacity to influence host QS and simultaneously enact its own lysis-lysogeny decision.

VqmA_Phage_ shares ∼43% amino acid sequence identity with VqmA_Vc_, and most of the key residues required for ligand and DNA binding are conserved (5,9). Thus, how VqmA_Phage_ can recognize two different promoters present in genomes from organisms in different kingdoms (viral and bacterial), while VqmA_Vc_ cannot, is not understood.

Here, we define the mechanism underlying this asymmetry. We show that VqmA selectivity for target promoters is driven by the DNA-binding domain (DBD) of the respective protein. We identify 6 key nucleotides within P*vqmR* and P*qtip* that contribute to VqmA promoter-binding selectivity, as exchanging these critical DNA sequences inverts the DNA-binding preferences of the two VqmA proteins. The 192^nd^ and 194^th^ residues in VqmA_Vc_ and VqmA_Phage_, respectively, within the helix-turn-helix (HTH) motifs, contribute to promoter-binding specificity. Isolation of VqmA_Phage_ mutants capable of activating *vqmR* expression but incapable of activating *qtip* expression revealed the residues G201, A202, E207, and M211 as crucial. These residues are either conserved (G201 and A202) or functionally conserved (E207 and M211) between VqmA_Phage_ and VqmA_Vc_, indicating that VqmA_Vc_ must possess an additional feature that prevents it from binding P*qtip* DNA. A mosaic VqmA_Vc_ protein containing the phage VqmA_Phage_ HTH motif along with the C-terminal 25 flanking VqmA_Phage_ residues was capable of binding P*qtip*. Thus, the two corresponding regions in VqmA_Vc_ must function in concert to prevent VqmA_Vc_ from binding to P*qtip*. Together, our analyses demonstrate how VqmA_Phage_, via its promiscuous DNA-binding activity, can control phage VP882 functions and drive host *V. cholerae* QS. Moreover, we discover why *V. cholerae* VqmA_Vc_ cannot do the reverse, as its DNA binding is strictly constrained to the host *V. cholerae* genome.

## RESULTS

### VqmA promoter-binding selectivity is conferred by the DNA-binding domain

VqmA proteins are composed of N-terminal Per-Arnt-Sim (PAS) domains responsible for binding the DPO AI and C-terminal DBDs containing HTH motifs (10). Both VqmA_Vc_ and VqmA_Phage_ bind DPO. By contrast, with respect to DNA binding, VqmA_Phage_ binds to P*qtip* and P*vqmR*, whereas VqmA_Vc_ only binds to P*vqmR* (5). We reasoned that this asymmetric DNA-binding pattern arises from differences in the DBDs (Supplementary Figure 1). To test this idea, we constructed chimeras in which we exchanged the VqmA_Vc_ and VqmA_Phage_ C-terminal domains to produce _Vc_N-C_Phage_ and _Phage_N-C_Vc_ proteins. We chose to make the junction at a residue near the C-terminal end of the PAS domain immediately following an amino acid stretch (GTIF) that is identical in both VqmA_Vc_ and VqmA_Phage_ (Supplementary Figure 1). We cloned *vqmA*_*Vc*_, *vqmA*_*Phage, Vc*_*N*-*C*_*Phage*_, and _*Phage*_*N*-*C*_*Vc*_ under an arabinose-inducible promoter and transformed each construct into recombinant Δ*tdh E. coli* harboring a P*vqmR-lux* or a P*qtip-lux* reporter. The Tdh enzyme is required for DPO biosynthesis, thus Δ*tdh E. coli* makes no DPO (3). Apo-VqmA displays basal transcriptional activity *in vivo* (9). Thus, using Δ*tdh E. coli* for these studies ensured that any transcriptional activity that occurred was exclusively a consequence of the DNA-binding capabilities of the chimeras and not ligand-binding-driven transcriptional activation of the chimeras. Consistent with our hypothesis, promoter activation by each chimera was determined by the protein from which the DBD originated: All four versions of VqmA activated P*vqmR-lux*, whereas only VqmA_Phage_ and _Vc_N-C_Phage_ activated P*qtip-lux* (Figure 1A and 1B, respectively). Next, we conjugated the four versions of VqmA into Δ*tdh* Δ*vqmA*_*Vc*_ *V. cholerae* lysogenized by a phage VP882 mutant in which the endogenous *vqmA*_*Phage*_ was inactive (VP882 *vqmA*_*Phage*_::Tn*5*). Thus, the only source of VqmA protein was that made from the plasmid. As expected, following arabinose-induction, only VqmA_Phage_ and _Vc_N-C_Phage_ activated *qtip* expression and induced host-cell lysis (Figure 1C).

**Figure 1.**
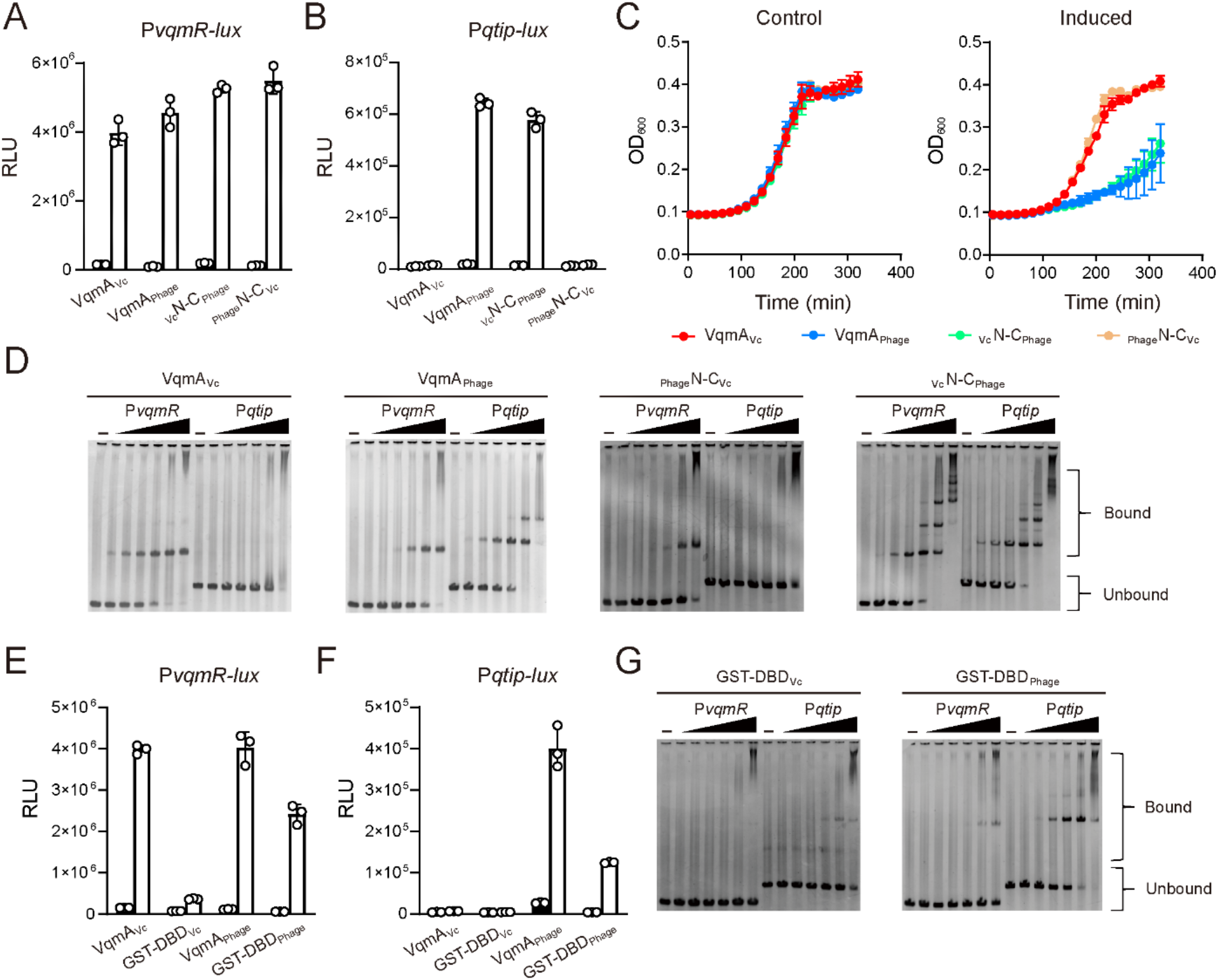
Promoter DNA-binding selectivity is conferred by the VqmA DBD. (A and B) Normalized reporter activity from Δ*tdh E. coli* harboring (A) P*vqmR-lux* or (B) P*qtip-lux* and arabinose-inducible VqmA_Vc_, VqmA_Phage, Vc_N-C_Phage_, or _Phage_N-C_Vc_. Black, no arabinose; white, 0.2% arabinose. Data are represented as mean ± SD (error bars) with *n=3* biological replicates. (C) Growth curves of the Δ*tdh* Δ*vqmA*_*Vc*_ *V. cholerae* harboring phage VP882 *vqmA*_*Phage*_::Tn*5* and arabinose-inducible VqmA_Vc_, VqmA_Phage, Vc_N-C_Phage_, or _Phage_N-C_Vc_ in medium lacking (Control) or containing 0.2% arabinose (Induced). (D) EMSAs showing binding of VqmA proteins to P*vqmR* and P*qtip* DNA. From left to right are, VqmA_Vc_, VqmA_Phage, Phage_N-C_Vc_, and _Vc_N-C_Phage_. 25 nM P*vqmR* or P*qtip* DNA was used in all EMSAs with no protein (designated -) or 2-fold serial dilutions of proteins. The lowest and highest protein (dimer) concentrations are 18.75 nM and 600 nM, respectively. (E and F) Normalized reporter activity from WT *E. coli* as in panels A and B harboring arabinose-inducible VqmA_Vc_, GST-DBD_Vc_, VqmA_Phage_, and GST-DBD_Phage_. (G) EMSAs showing binding of GST-DBD_Vc_, and GST-DBD_Phage_ to P*vqmR* and P*qtip* DNA. Probe and protein concentrations as in panel D.

We verified the above findings *in vitro* using electrophoretic mobility shift assays (EMSAs). Consistent with the cell-based assays, the purified VqmA_Vc_, VqmA_Phage, Vc_N-C_Phage_, and _Phage_N-C_Vc_ proteins shifted P*vqmR* DNA, whereas only the VqmA_Phage_ and _Vc_N-C_Phage_ proteins shifted P*qtip* DNA (Figure 1D). Assessing the ratios of bound to total DNA across varying protein concentrations allowed us to calculate the relative binding affinities (EC_50_) of the VqmA proteins for P*vqmR* and P*qtip* DNA (Supplementary Figure 2A). _Phage_N-C_Vc_, like VqmA_Vc_, only bound P*vqmR*, but with a ∼7-fold lower affinity. Consistent with our previous findings, VqmA_Phage_ bound P*qtip* about 3-fold more strongly than it bound P*vqmR* (5). By contrast, _Vc_N-C_Phage_ showed a modest increase in its preference for P*qtip* relative to that for P*vqmR*, with binding to both promoters at a level similar to that with which VqmA_Phage_ bound P*qtip*. Indeed, in agreement with our EC_50_ measurements, when P*qtip* and P*vqmR* DNA were supplied at equimolar concentrations in a competitive DNA-binding assay, lower amounts of VqmA_Phage_ and _Vc_N-C_Phage_ were required to shift P*qtip* DNA than to shift P*vqmR* DNA (Supplementary Figure 2B). In conclusion and in agreement with our *in vivo* results, the respective DBD of each purified VqmA protein drives promoter selectively.

The VqmA_Vc_ PAS domain is responsible for sensing DPO and for dimerization (9,11). To eliminate the possibility that the PAS domain also plays a role in promoter-binding selectivity, we assayed the VqmA_Vc_ and VqmA_Phage_ DBDs lacking their partner PAS domains (DBD_Vc_ and DBD_Phage_, respectively) for activation of P*vqmR-lux* and P*qtip-lux*. Neither DBD activated transcription (Supplementary Figure 3A and 3B, respectively). Likewise, EMSA analyses showed that neither DBD bound either promoter (Supplementary Figure 3C). Gel filtration analyses indicated that the DBD proteins purified as monomers (Supplementary Figure 3D), suggesting that, indeed, the DBDs were unable to dimerize in the absence of their partner PAS domains.

Transcriptional activity driven by HTH-containing proteins typically depends on dimer formation. Soluble glutathione S-transferase (GST) spontaneously forms a homodimer (12), and so GST can be employed as a substitute for native dimerization domains of proteins (13). Thus, to examine the VqmA requirement for dimerization, we fused GST to the N-terminus of each VqmA DBD to yield recombinant GST-DBD_Vc_ and GST-DBD_Phage_ and we tested whether DNA-binding function was restored. Indeed, the GST-DBD proteins purified as dimers (Supplementary Figure 3D). P*vqmR-lux* and P*qtip-lux* expression analyses revealed that the DBDs, when fused to GST, regained function, with the caveat that the GST-DBD_Vc_ exhibited 10-fold reduced activity compared to WT VqmA_Vc_ (Figure 1E). Importantly, the DNA-binding preferences mimicked those of the full-length proteins: GST-DBD_Phage_ activated both P*vqmR-lux* and P*qtip-lux*, whereas GST-DBD_Vc_ only activated P*vqmR-lux* (Figure 1E and 1F). Companion EMSA analyses showed that GST-DBD_Phage_ bound P*qtip* ∼5-fold more strongly than it bound P*vqmR*, whereas GST-DBD_Vc_ showed almost no binding to P*vqmR* and, unexpectedly, some weak binding could be detected to the P*qtip* DNA (Figure 1G). We confirmed that purified GST alone did not bind either P*vqmR* or P*qtip* (Supplementary Figure 3E). Given that the GST-DBD_Vc_ driven activation of P*qtip-lux* was undetectable *in vivo* (Figure 1F), we presume that the observed *in vitro* GST-DBD_Vc_ binding to P*qtip* DNA is a consequence of the simplified context in which the EMSA is performed. Likely, the DNA:VqmA ratio in the EMSA is far higher than in cells, which, in the case of GST-DBD_Vc_, fosters modest non-specific DNA binding. Taken together, our results show that dimerization is necessary for VqmA DNA-binding function, and that VqmA promoter-binding selectivity is governed exclusively by the DBD.

### VqmA DNA-binding preferences can be inverted by exchanging key DNA sequences in PvqmR and Pqtip

To study the VqmA promoter-binding asymmetry from the aspect of the DNA, our next goal was to identify the critical DNA sequence within P*qtip* that prevents VqmA_Vc_ from binding. In the phage VP882 genome, P*qtip* resides between *vqmA*_*Phage*_ and *qtip* and VqmA_Phage_ activates its own and *qtip* expression, suggesting that VqmA_Phage_ binding may involve both DNA strands. Similarly, VqmA_Vc_ has been shown to interact with both strands of the P*vqmR* (11). Thus, in each case, both DNA strands need to be considered (Figure 2A). Previous work revealed that the critical region in P*vqmR* required for VqmA_Vc_ binding is -AGGGGGGATTTCCCCCCT-(2,11). The corresponding fragment from P*qtip*, but on the opposite DNA strand, -TAGGGGGAAAAATACCCT-, possesses ∼56% sequence identity to this region suggesting it could be the key stretch of DNA that drives VqmA_Phage_ promoter selection. The highest divergence in the two promoters is in the central 6 nucleotides: “-AAAATA-” in P*qtip* and “-TTTCCC-” in P*vqmR*. We synthesized DNA probes in which we exchanged the “-AAAATA-” in P*qtip* with “-TTTCCC-” from P*vqmR* and tested VqmA_Vc_ and VqmA_Phage_ binding by EMSA analysis. We call these probes P*vqmR*\* and P*qtip*\*, respectively. Indeed, promoter DNA-binding specificity was exchanged: VqmA_Vc_ only shifted P*qtip\**, and VqmA_Phage_ bound to P*vqmR\** twice as strongly as it bound to P*qtip\**, showing the opposite preference for the two synthetic promoters compared to the native promoters (Figure 2B). P*vqmR*-lux* and P*qtip*-lux* transcriptional fusions mimicked the EMSA results: VqmA_Vc_ only activated expression of P*qtip*-lux*, whereas VqmA_Phage_ activated expression of P*vqmR*-lux* and P*qtip\**-*lux* (Figure 2C and 2D). Thus, this 6-nucleotide stretch is the key sequence that determines the DNA-binding specificity for the two VqmA proteins. Moreover, the presence of the -AAAATA-nucleotide sequence in P*qtip* is sufficient to prevent VqmA_Vc_ from binding.

**Figure 2.**
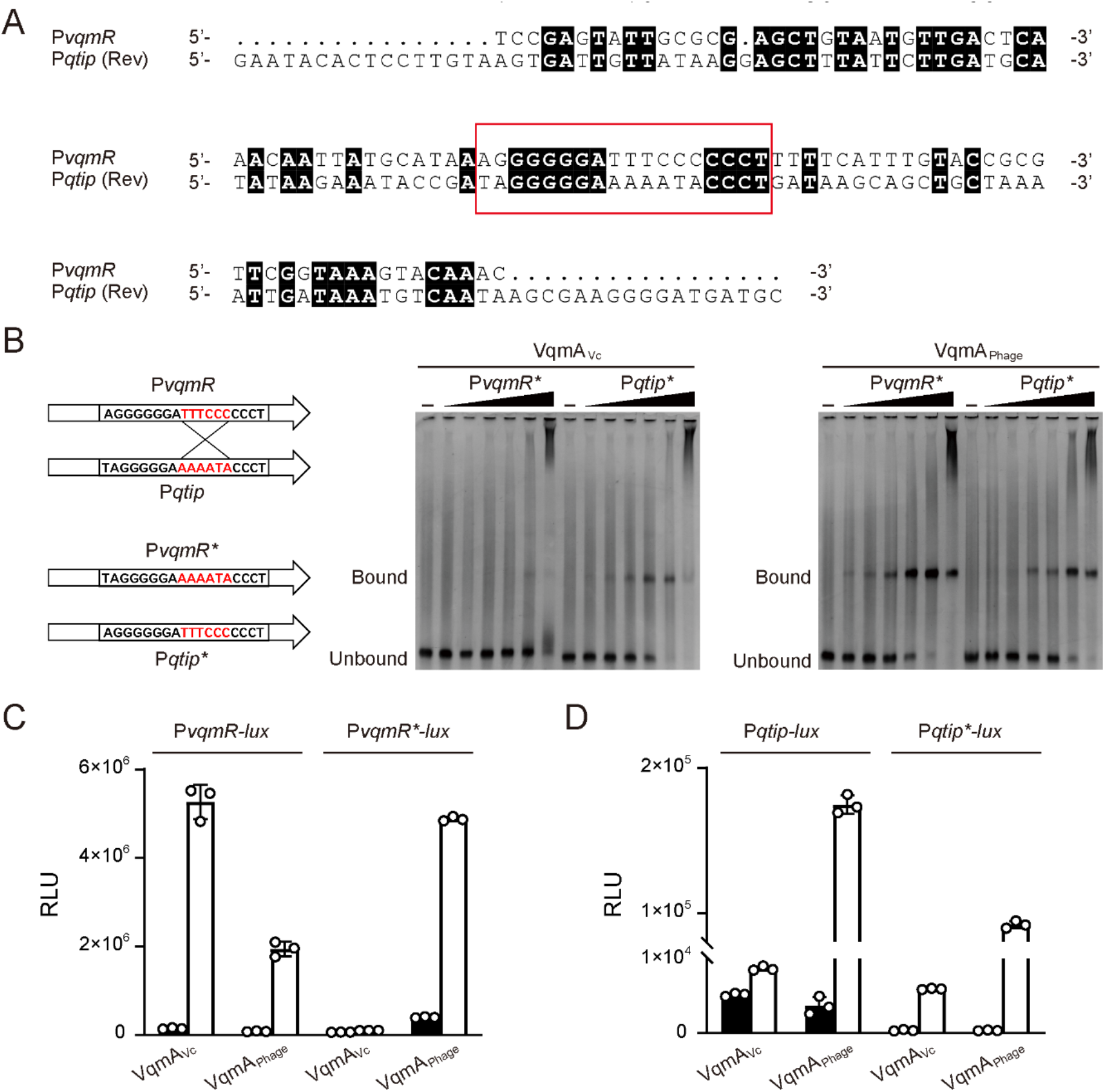
Promoter selectivity is reversed by exchanging key nucleotide fragments. (A) DNA sequence alignment (ClustalW) of P*vqmR* and P*qtip*. The reverse strand of P*qtip* is shown. Identical nucleotides are designated with black shading. (B) EMSAs showing binding of the designated VqmA proteins to P*vqmR*\* and P*qtip*\* DNA. The cartoon at the left illustrates the key sequences exchanged in the probes. Probe and protein concentrations as in Figure 1D. (C) Normalized reporter activity from Δ*tdh E. coli* harboring P*vqmR-lux* or P*vqmR*-lux* and arabinose-inducible VqmA_Vc_ or VqmA_Phage_. Black, no arabinose; white, 0.2% arabinose. Data are represented as mean ± SD (error bars) with *n=3* biological replicates. (D) As in C for P*qtip-lux* or P*qtip*-lux*.

### Protein sequence-guided mutagenesis reveals that residue E194 in phage VP882 VqmA_Phage_ and the equivalent A192 residue in V. cholerae VqmA_Vc_ contribute to specificity for Pqtip

We considered two possible mechanisms that could underpin the asymmetric VqmA DNA-binding patterns: phage VP882 VqmA_Phage_ could possess a feature that relaxes its DNA-binding specificity, and/or *V. cholerae* VqmA_Vc_ could possess a feature that restricts its DNA-binding ability. To distinguish between these possibilities, we first probed which residues drive VqmA_Phage_ interactions with P*qtip* but do not contribute to interactions with P*vqmR*. To do this, we performed site-directed mutagenesis of VqmA_Phage_ with the goal of identifying mutants that fail to bind P*qtip* but retain binding to P*vqmR*. Charged residues in HTH motifs typically mediate interactions between VqmA-type transcription factors and DNA, and indeed, both VqmA HTHs are enriched in positively-charged amino acids (9,11). Sequence alignment of the HTHs in VqmA_Phage_ and VqmA_Vc_ revealed four obvious differences in charged residues that could underlie the DNA-binding asymmetry between the two proteins (Supplementary Figure 1). We mutated those residues in VqmA_Phage_ to the corresponding VqmA_Vc_ residues. The changes are: VqmA_Phage_^K176Q^, VqmA_Phage_^R184I^, VqmA_Phage_^I193E^, and VqmA_Phage_^E194A^. To test the combined effect of these mutations on VqmA_Phage_ DNA-binding function, we also constructed the quadruple VqmA_Phage_^K176Q, R184I, I193E, E194A^mutant. VqmA_Phage_^K176Q^, VqmA_Phage_^R184I^, VqmA_Phage_^I193E^retained the ability to induce phage lysis showing that *in vivo* binding to P*qtip* was not eliminated (Figure 3A). VqmA_Phage_^E194A^induced only low-level cell lysis suggesting that, while binding to P*qtip* is not eliminated, it is compromised (Figure 3A). Analysis of P*vqmR-lux* and P*qtip-lux* expression revealed that all four VqmA_Phage_ single point mutants possessed WT P*vqmR-lux* activity and ∼2-7-fold reductions in P*qtip-lux* activity with VqmA_Phage_^E194A^being the least active (Figure 3B and 3C, respectively). The quadruple mutant was unable to induce phage lysis in a *V. cholerae* lysogen and it did not activate P*vqmR-lux* or P*qtip-lux* expression showing it is defective in binding to both promoters (Figure 3A-3C). Western blot analysis demonstrated that all of the VqmA_Phage_ variants were produced at levels similar to WT in both *V. cholerae* and *E. coli* (Supplementary Figure 4A). Thus, our results indicate that, among these charged residues, only the VqmA_Phage_ residue E194 in the HTH motif plays a role in VqmA_Phage_ selection of P*qtip*.

**Figure 3.**
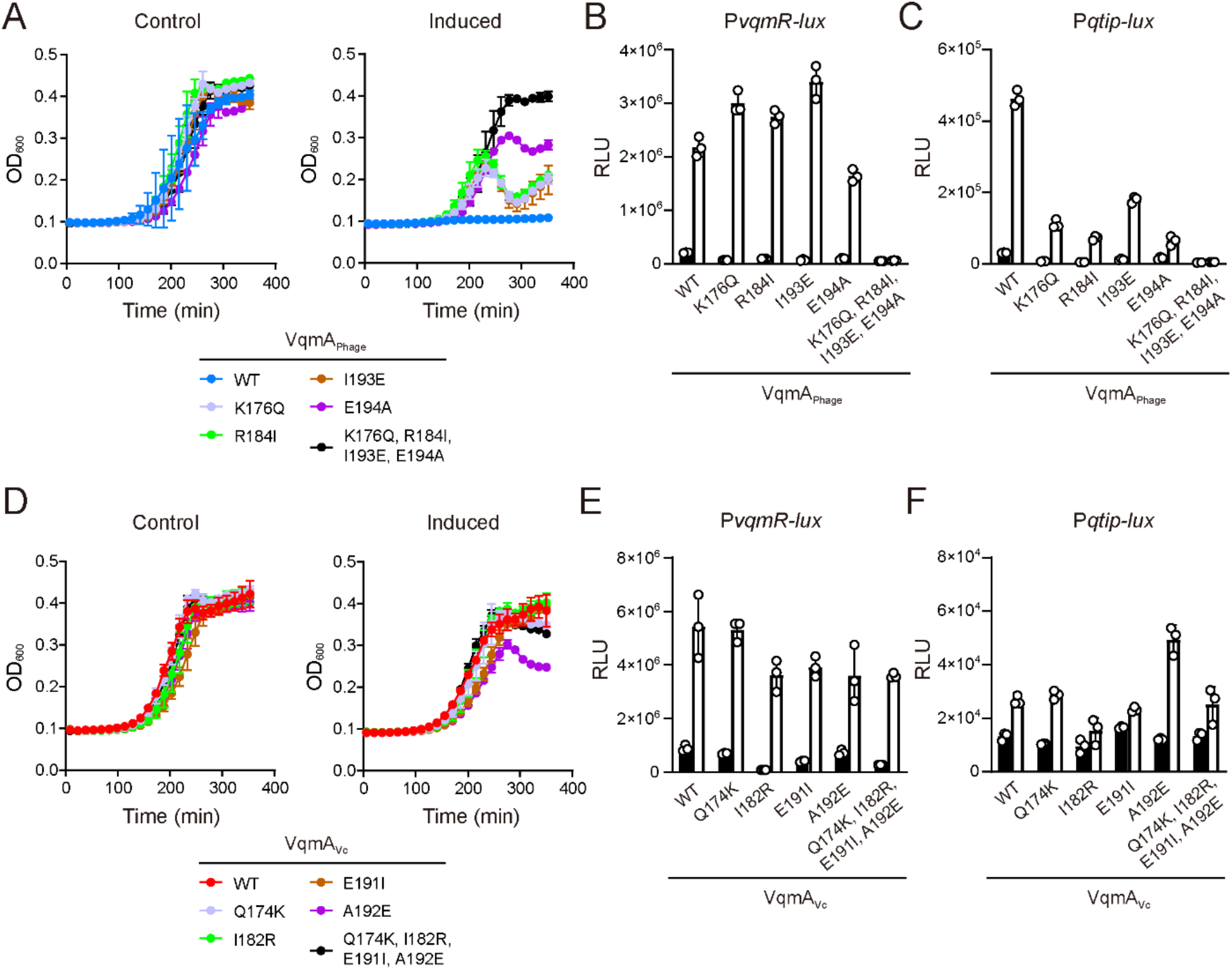
VqmA_Phage_ residue E194 and the corresponding VqmA_Vc_ residue A192 contribute to specificity for binding to P*qtip*. (A) Growth curves of Δ*tdh* Δ*vqmA*_*Vc*_ *V. cholerae* harboring phage VP882 *vqmA*_*Phage*_::Tn*5* and the indicated 3xFLAG-VqmA_Phage_ alleles in medium lacking (Control) or containing 0.2% arabinose (Induced). (B and C) Normalized reporter activity from Δ*tdh E. coli* harboring (B) P*vqmR-lux* or (C) P*qtip-lux* and the indicated arabinose-inducible 3xFLAG-VqmA_Phage_ alleles. Black, no arabinose; white, 0.2% arabinose. Data are represented as mean ± SD (error bars) with *n=3* biological replicates. (D) Growth curves of Δ*tdh* Δ*vqmA*_*Vc*_ *V. cholerae* lysogens harboring the indicated 3xFLAG-VqmA_Vc_ alleles in medium lacking (Control) or containing 0.2% arabinose (Induced). (E and F) Normalized reporter activity from Δ*tdh E. coli* harboring (E) P*vqmR-lux* or (F) P*qtip-lux* and the indicated arabinose-inducible 3xFLAG-VqmA_Vc_ alleles. Black, no arabinose; white, 0.2% arabinose. Data are represented as mean ± SD (error bars) with *n=3* biological replicates.

While the residues we mutated in the phage VP882 VqmA_Phage_ HTH motif do not dramatically perturb site-specific recognition of P*qtip*, the corresponding residues in the *V. cholerae* VqmA_Vc_ HTH motif could nonetheless restrict its capacity to bind P*qtip*. Therefore, we also mutated the analogous VqmA_Vc_ residues to the corresponding VqmA_Phage_ residues. We made: VqmA_Vc_ ^Q174K^, VqmA_Vc_ ^I182R^, VqmA_Vc_ ^E191I^, VqmA_Vc_^A192E^ and VqmA_Vc_ ^Q174K, I182R, E191I, A192E^. Here, our goal was to test whether the variants gained the ability to bind P*qtip*. Only VqmA_Vc_^A192E^ induced a modest level of lysis in the *V. cholerae* lysogen, whereas all other VqmA_Vc_ variants failed to do so (Figure 3D). All of the VqmA_Vc_ variants drove the WT level of P*vqmR-lux* activity (Figure 3E). VqmA_Vc_^A192E^ generated low but detectable P*qtip-lux* expression, while the other VqmA_Vc_ variants did not (Figure 3F). The VqmA_Vc_ variants were produced at similar levels to WT VqmA_Vc_ in *V. cholerae* and *E. coli* (Supplementary Figure 4B). We conclude that, among the tested residues, only A192 plays a role in preventing VqmA_Vc_ from binding P*qtip*.

Our mutagenesis analyses for VqmA_Vc_ are consistent with our analyses for VqmA_Phage_: the residue at the 192^nd^ position in *V. cholerae* VqmA_Vc_ and the analogous residue at the 194^th^ position in phage VP882 VqmA_Phage_ contribute to selection of P*qtip*. However, given that the A192E substitution in VqmA_Vc_ results in only partial activation of P*qtip* expression, and the E194A substitution in VqmA_Phage_ results in only partial loss of activation of P*qtip*, the E194 residue in VqmA_Phage_ cannot be the sole amino acid responsible for the preference VqmA_Phage_ shows for P*qtip*. Rather, additional residues in VqmA_Phage_ must participate in conferring specificity.

### Random mutagenesis of the VqmA_Phage_ DBD reveals that residues G201, A202, E207, and M211 are required for VqmA_Phage_ to bind Pqtip but are dispensable for binding PvqmR

Our protein sequence-guided approach did not reveal the primary mechanism underlying promoter-binding specificity for either of the VqmA proteins. We therefore performed a genetic screen to forward our goal of identifying phage VP882 VqmA_Phage_ mutants that fail to bind P*qtip* but retain the ability to bind P*vqmR*. We constructed a library of random mutations in the *vqmA*_*Phage*_ DBD, cloned them into a plasmid under an arabinose-inducible promoter, and introduced them into Δ*tdh* Δ*vqmA*_*Vc*_ *V. cholerae* harboring P*vqmR*-*lux* on the chromosome and lysogenized by phage VP882 harboring inactive *vqmA*_*Phage*_ (*vqmA*_*Phage*_::Tn*5*). The logic of the screen is as follows: When propagated on agar plates supplemented with arabinose, *V. cholerae* exconjugants harboring *vqmA*_*Phage*_ alleles possessing reasonable P*qtip*-binding activity will lyse because those VqmA_Phage_ proteins will bind P*qtip* on the phage VP882 genome and launch the phage lytic cascade (Supplementary Figure 5). Such exconjugants will die and thus be eliminated from the screen. Exconjugants that survive but carry *vqmA*_*Phage*_ null alleles will produce no light because those VqmA_Phage_ proteins will fail to bind P*vqmR-lux*, so they also can be eliminated from the screen. The *vqmA*_*Phage*_ alleles of interest to us are those that are maintained in surviving exconjugants (because they encode proteins that cannot bind P*qtip*) and produce light (because they encode proteins that can bind P*vqmR-lux*).

Our screen yielded the following mutants: VqmA_Phage_^G201D^, VqmA_**Phage**_^**G201R**^, VqmA_Phage_ ^A202V^, VqmA_Phage_^E207K^, VqmA_Phage_^E207V^, and VqmA_Phage_^M211K^ (Figure 4A). To verify that these VqmA_Phage_ mutants were indeed defective in binding P*qtip*, we individually transformed them into Δ*tdh E. coli* carrying the P*qtip-lux* reporter or the P*vqmR-lux* reporter and measured light production. All variants retained WT capability to activate P*vqmR-lux*, but they did not harbor WT capability to activate P*qtip-lux* expression (>10-fold reductions in activity) (Figure 4B and 4C, respectively). Thus, any residual P*qtip* binding by these mutant VqmA_Phage_ proteins is insufficient to induce host-cell lysis in the phage VP882 lysogen (Figure 4A). We verified that the VqmA_Phage_ variants are produced at the same level as WT VqmA_Phage_ in *V. cholerae* and *E. coli* (Supplementary Figure 4C). According to the protein sequence alignment, VqmA_Phage_ residues (175-200) corresponding to positions 173-198 in VqmA_Vc_ comprise the VqmA_Phage_ HTH motif (Supplementary Figure 1). Thus, the residues identified in the mutagenesis (G201, A202, E207, and M211) are located C-terminal to the VqmA_Phage_ HTH motif. Mapping the analogous *V. cholerae* VqmA_Vc_ residues (G199, A200, Q205, and L209) to the DPO-VqmA_Vc_-DNA structure also shows that all of these residues cluster in a flexible loop region and helix adjacent to, but distinct from the HTH motif that directly contacts DNA (Figure 4D; Supplementary Figure 1). Surprisingly, the residues identified in the VqmA_Phage_ mutagenesis are either identical (VqmA_Phage_ G201 and A202 versus VqmA_Vc_ G199 and A200) or similar (VqmA_Phage_ E207 and M211 versus VqmA_Vc_ Q205 and L209) between VqmA_Phage_ and VqmA_Vc_. To test whether possession of the similar residues is sufficient to confer DNA-binding specificity for P*qtip*, we constructed VqmA_Vc_ ^Q205E^ and VqmA_Vc_^L209M^ and tested their DNA-binding functions as above. VqmA_Vc_ ^Q205E^ and VqmA_Vc_ ^L209M^, like WT VqmA_Vc_, activated P*vqmR-lux* but failed to activate P*qtip-lux* (Supplementary Figure 6A and 6B, respectively). We make the following four conclusions from these findings: 1) There are four residues (G201, A202, E207, and M211) required for VqmA_Phage_ to recognize P*qtip* DNA. 2) Because the VqmA_Phage_ G201D, G201R, A202V, E207K, E207V, and M211K variants exhibit WT binding to P*vqmR*, the four residues G201, A202, E207, and M211 must not play any role in P*vqmR* recognition. 3) Both VqmA_Phage_ and VqmA_Vc_ possess the capacity to bind P*qtip* DNA. 4) Because VqmA_Vc_ in fact does not bind P*qtip*, VqmA_Vc_ must possess an additional feature that prevents that from occurring.

**Figure 4.**
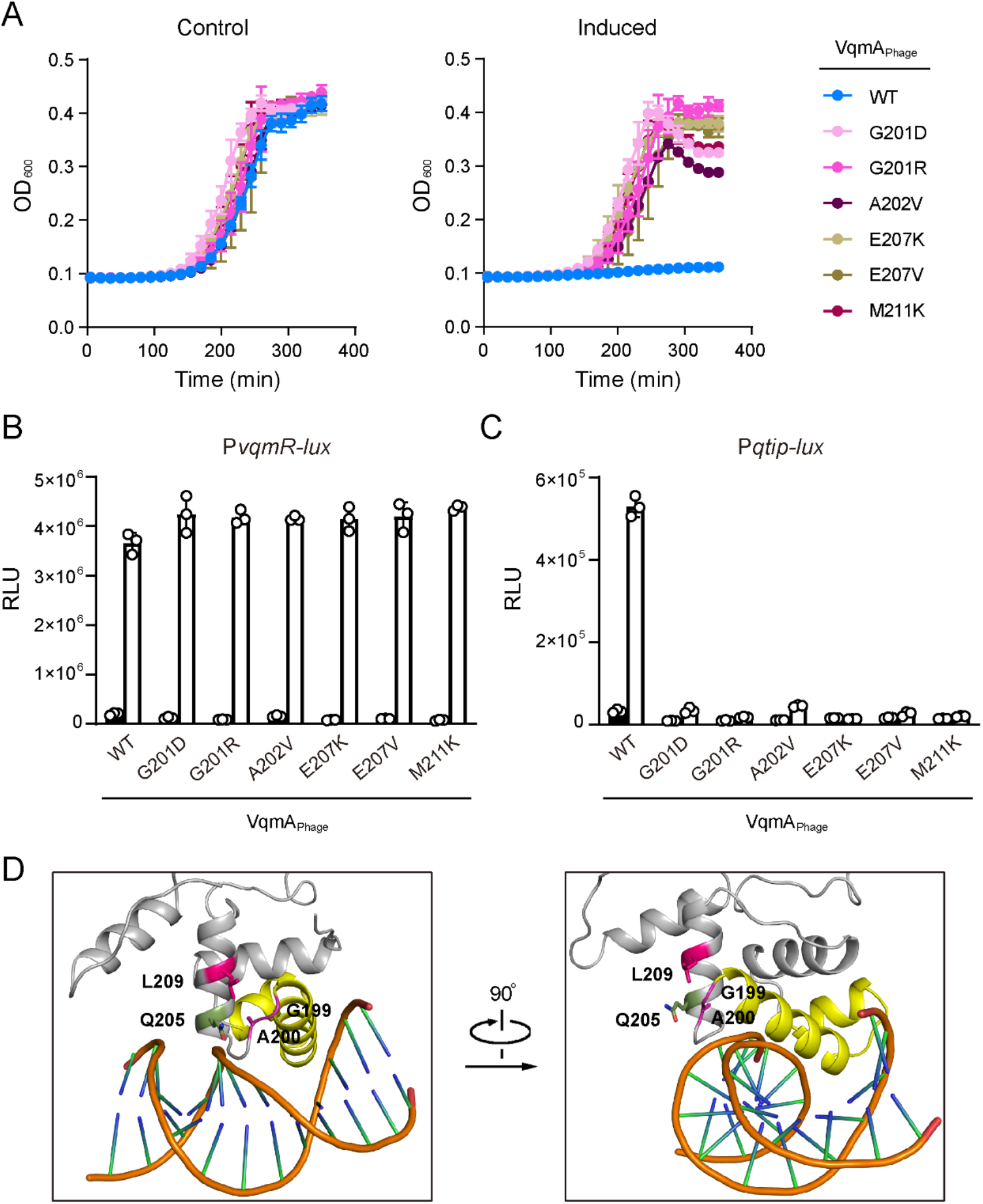
VqmA_Phage_ residues G201, A202, E207, and M211 are required for binding to P*qtip*. (A) Growth curves of Δ*tdh* Δ*vqmA*_*Vc*_ *V. cholerae* lysogens harboring the indicated VqmA_Phage_ alleles in medium lacking (Control) or containing 0.2% arabinose (Induced). (B and C) Normalized reporter activity from Δ*tdh E. coli* harboring (B) P*vqmR-lux* or (C) P*qtip-lux* and the indicated arabinose-inducible 3xFLAG-VqmA_Phage_ alleles. Black, no arabinose; white 0.2% arabinose. Data are represented as mean ± SD (error bars) with *n=3* biological replicates. (D) Close up views of the DBD from the crystal structure of DPO-VqmA_Vc_ bound to DNA (PDB: 6ide, protein in gray with the HTH motif in yellow, and the DNA in orange). The color scheme for VqmA_Vc_ residues G199, A200, Q205, and L209 mirrors that used in panel A.

### The restrictive element that prevents VqmA_Vc_ from binding Pqtip is located in its HTH motif and the adjacent C-terminal region of 25 residues

To test the hypothesis that a feature in the VqmA_Vc_ DBD restricts its DNA-binding capacity for P*vqmR*, we performed a genetic screen aimed at identifying VqmA_Vc_ mutants capable of activating P*qtip-lux* expression. To do this, we constructed a library of random *vqmA*_*Vc*_ DBD alleles and cloned them into a plasmid under an arabinose-inducible promoter. The library was transformed into the Δ*tdh E. coli* strain harboring the P*qtip-lux* reporter and transformants were propagated on plates containing arabinose. We screened for transformants that produced light indicating that they contained VqmA_Vc_ proteins that activated P*qtip-lux*. This strategy yielded no such transformants. Several possibilities could explain our result: we did not screen sufficient numbers of mutants, the mutagenesis did not yield the crucial change, or no alteration of a single residue can enable VqmA_Vc_ binding to P*qtip*.

We expanded our search for the DNA-binding restrictive element present in VqmA_Vc_ by assessing whether a particular region in the VqmA_Vc_ DBD constrains promoter binding to P*vqmR*. To do this, we constructed five VqmA_Vc_ mosaic proteins by replacing ∼20-30 residues in the *V. cholerae* VqmA_Vc_ DBD with the corresponding residues from the phage VP882 VqmA_Phage_ DBD. We call these proteins VqmA_Vc_^*126-149^, VqmA_Vc_ ^*150-170^, VqmA_Vc_ ^*171-199^, VqmA_Vc_ ^*200-224^, and VqmA_Vc_ ^*225-246^ (see Supplementary Figure 1 for relevant protein segments). Each superscript denotes the VqmA_Vc_ amino acid residues that have been replaced by the corresponding residues from VqmA_Phage_. In all the mosaics, either the intact VqmA_Vc_ HTH or the intact VqmA_Phage_ HTH was present. For reference, the VqmA_Vc_ HTH motif consists of residues 173 to 198 and the VqmA_Phage_ HTH spans residues 175 to 200. We tested the mosaic VqmA_Vc_ proteins for activation of the P*vqmR-lux* and P*qtip-lux* reporters. The DNA specificity of all the VqmA_Vc_ mosaics mimicked WT VqmA_Vc_ as P*vqmR-lux* was expressed but P*qtip-lux* was not (Figure 5A and 5B, respectively). We confirmed that the mosaic VqmA_Vc_ proteins are expressed at levels similar to WT VqmA_Vc_ (Supplementary Figure 7A). Our results suggest that the feature that prevents *V. cholerae* VqmA_Vc_ from binding to P*qtip* is larger than the regions delineated by any of the VqmA_Vc_ mosaics, or it could be that multiple patches in the VqmA_Vc_ DBD that are not contiguous in amino acid sequence are responsible.

**Figure 5.**
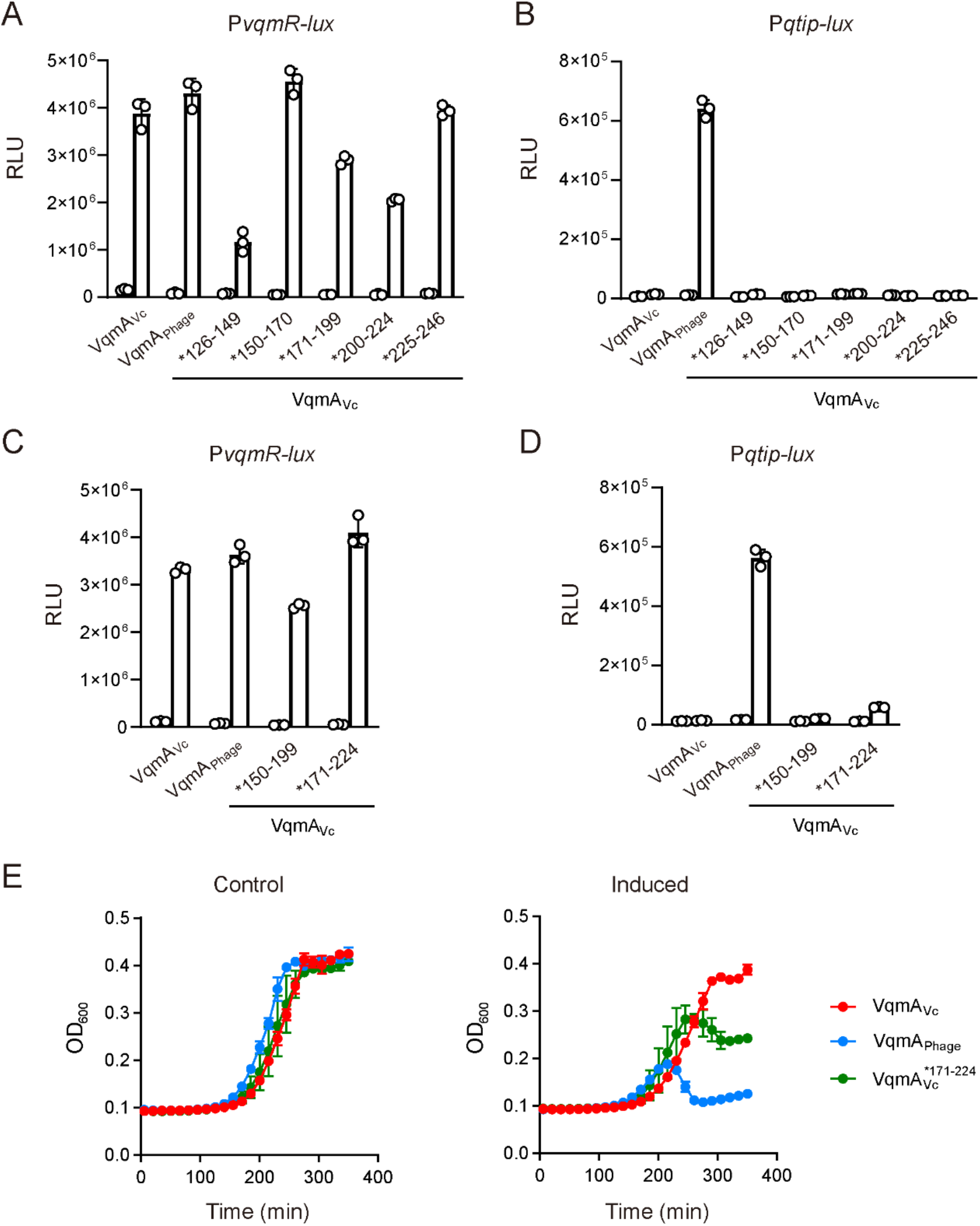
The VqmA_Vc_ HTH motif and the immediate C-terminal 25 residues, together, constrain binding to P*vqmR*. (A-D) Normalized reporter activity from Δ*tdh E. coli* harboring (A and C) P*vqmR-lux* or (B and D) P*qtip-lux* and arabinose-inducible VqmA_Vc_, VqmA_Phage_, or the indicated VqmA_Vc_ allele. Data are represented as mean ± SD (error bars) with *n=3* biological replicates. Black, no arabinose; white, 0.2% arabinose. (E) Growth curves of Δ*tdh* Δ*vqmA*_*Vc*_ *V. cholerae* lysogens harboring VqmA_Vc_, VqmA_Phage_, or VqmA_Vc_ ^*171-224^ in medium lacking (Control) or containing 0.2% arabinose (Induced).

Pinpointing non-contiguous regions that could, together, contain the VqmA_Vc_ restrictive element is challenging. However, testing for a larger contiguous expanse that could contain the putative restrictive element is straightforward. Thus, we constructed two additional *V. cholerae* VqmA_Vc_ mosaic proteins. In one construct, called VqmA_Vc_ ^*150-199^, we introduced the VqmA_Phage_ HTH along with the immediate N-terminal 25 amino acids in place of the corresponding VqmA_Vc_ region. Second, in a construct called VqmA_Vc_^*171-224^, we introduced the VqmA_Phage_ HTH together with the immediate C-terminal 25 amino acid stretch in place of that VqmA_Vc_ region. VqmA_Vc_^*150-199^and VqmA_Vc_^*171-224 >^ activated P*vqmR-lux* to approximately WT levels, whereas only VqmA_Vc_^*171-224^activated P*qtip-lux*, albeit weakly (Figure 5C and 5D, respectively). Consistent with this result, VqmA_Vc_ ^*171-224^ induced partial lysis in the *V. cholerae* phage VP882 lysogen (Figure 5E). VqmA_Vc_ ^*171-224^ was produced at levels similar to WT VqmA_Vc_, eliminating the possibility that the observed binding to P*qtip* was a consequence of overexpression (Supplemental Figure 7B). We conclude that the region encompassing both the HTH motif and the C-terminal 25 residues are required to restrict the VqmA_Vc_ DBD from binding P*qtip*.

## DISCUSSION

The DPO-VqmA QS AI-receptor pair controls lifestyle transitions in the pathogen *V. cholerae* and in the vibriophage VP882. Here, we studied the DNA-binding function of VqmA. VqmA proteins are cytoplasmic transcription factors composed of N-terminal PAS domains responsible for binding the DPO ligand and C-terminal DBDs containing HTH motifs. Most of the key residues required for binding the DPO ligand and for binding to P*vqmR* DNA are conserved between the two VqmA proteins. Indeed, both VqmA_Vc_ and VqmA_Phage_ bind DPO and activate transcription of *vqmR*. By contrast, only VqmA_Phage_ activates the phage gene *qtip*. Here, we investigated this asymmetric DNA-binding pattern. Our work shows that, in both proteins, the DBD determines promoter recognition. We have previously shown that DPO binding enhances VqmA transcriptional activity (9). This earlier work, together with our present results, suggest a model in which the PAS domain specifies DNA-binding affinity (between the apo- and holo-states), and the DBD specifies DNA-binding selectivity.

The main goal of the present work was to discover features of the VqmA proteins that confer specificity in transcriptional activity. Our hypothesis was that either phage VP882 VqmA_Phage_ could possess a feature that relaxes its DNA-binding specificity, and/or *V. cholerae* VqmA_Vc_ could possess a feature that restricts its DNA-binding capability. Our genetic analyses support the latter possibility and suggest that the VqmA_Vc_ DBD harbors elements that prevent it from binding P*qtip*. This hypothesis stems from our finding that the residues G201, A202, E207, and M211 are crucial for VqmA_Phage_ recognition of P*qtip*. These residues are conserved between VqmA_Vc_ and VqmA_Phage_. Specifically, in VqmA_Vc_ they are: G199, A200, Q205, and L209, respectively. More broadly, sequence alignments of VqmA proteins among *Vibrios* reveal that the residue at the 207^th^ position in VqmA_Phage_ (205^th^ position in VqmA_Vc_) is most frequently either a Glu or a Gln (5). Similarly, the residue at the 211^th^ position in VqmA_Phage_ (209^th^ position in VqmA_Vc_) is commonly a hydrophobic residue, like Met, Leu, Ile, or Val. Thus, E207 and M211 are not unique to VqmA_Phage_, but rather occur in most VqmA proteins. We propose that because the key residues for P*qtip* binding are conserved in VqmA_Phage_, VqmA_Vc_, and other *Vibrio* VqmA proteins, VqmA_Vc_ could bind P*qtip*, but is restricted from doing so by additional features in its DBD.

In the case of VqmA_Phage_, the residues G201, A202, E207, and M211 identified in our mutagenesis screen as necessary for P*qtip* binding are, surprisingly, not in the HTH motif, nor do the corresponding VqmA_Vc_ residues make direct contacts with DNA in the DPO-VqmA_Vc_-DNA (P*vqmR*) crystal structure (Figure 4D). Thus, we wonder how the G201, A202, E207, and M211 residues could govern recognition of P*qtip*. Our *in vivo* analyses showed that substitutions in VqmA_Phage_ at these residues enable activation of *vqmR* expression to WT levels, whereas only residual activation of *qtip* expression occurs (Figure 4A-C). Surprisingly, the purified VqmA_Phage_ mutant proteins maintained some capability to bind P*qtip in vitro*. A representative experiment using the VqmA_Phage_^G201D^protein is shown in Supplementary Figure 8.

We consider several possibilities to explain our findings:

First, the VqmA_Phage_ G201, A202, E207, and M211 residues could mediate interactions with an additional bacterial factor involved in transcription. Importantly, the failure of these VqmA_Phage_ variants to activate P*qtip* expression in *V. cholerae* lysogens also occurred in *E. coli*, eliminating the possibility that these residues interact with a phage-specific or *Vibrio*-specific factor. Rather, these residues could be important for coordinating interactions with a conserved factor, such as RNA polymerase. If so, these mutant VqmA_Phage_ proteins, while capable of binding promoter DNA, are incapable of activating transcription. This situation would be analogous to the positive control mutants of the lambda phage cI repressor (cI_lambda_). So called pc mutants bind DNA and exhibit repressor activity, but are deficient in positive transcriptional regulation due to the inability of the mutant cI_lambda_ proteins to productively interact with RNA polymerase (14,15). In our case, the VqmA_Phage_ mutants maintain the capacity to activate *vqmR* expression so they must successfully interact with RNA polymerase at least at P*vqmR*. For this reason, we consider it unlikely that these VqmA_Phage_ mutants are analogous to lambda pc mutants.

Second, a global transcriptional regulator could be involved that is present in both *V. cholerae* and *E. coli*. One candidate is the histone-like nucleoid structuring protein (H-NS) that functions as a universal repressor of transcription (16). In *Vibrio harveyi*, the QS master regulator, LuxR, displaces H-NS at promoter DNA to activate expression of QS-controlled genes (17). Perhaps, the VqmA_Phage_ G201, A202, E207, and M211 mutants cannot successfully compete with H-NS for binding at P*qtip in vivo*, whereas in an EMSA assay, since H-NS is not present, binding to P*qtip* DNA occurs. Any potential role for H-NS in transcriptional regulation of *vqmR* or *qtip* has not been investigated.

Third, the binding of the VqmA_Phage_ G201, A202, E207, and M211 mutants to P*qtip in vitro*, while demonstrating loss of activity *in vivo*, could be a consequence of the unnaturally high DNA: VqmA_Phage_ stoichiometry in the EMSA, similar to what we observed for the GST-DBD_Vc_ construct (Figure 1G). Thus, the EMSA is not sufficiently sensitive to distinguish between the strength of DNA binding of WT VqmA_Phage_ and the residual binding by the VqmA_Phage_ G201, A202, E207, and M211 mutants. If this is the case, we propose that VqmA_Phage_ G201, A202, E207, and M211 could play allosteric roles in correctly positioning the VqmA_Phage_ HTH for proper contact with particular DNA nucleotides. Here, we compare this possibility to how site-specific recognition is accomplished by cI_lambda_. Genetic and biochemical studies revealed that residues outside of the cI_lambda_ HTH motif are crucial for site-specific DNA recognition (18–22). The crystal structure of the cI_lambda_ repressor bound to DNA shows that charged residues adjacent to those in the HTH interact with the DNA sugar phosphate backbone (23). Additionally, the N-terminal arm of cI_lambda_ wraps around the DNA and makes contacts on the backside of the helix (23). It is presumed that the backbone contacts function to position the HTH residues to contact specific DNA nucleotides. Thus, while the VqmA_Phage_ residues that we identified as important for P*qtip* recognition (G201, A202, E207, and M211) do not function perfectly analogously to those in cI_lambda_ because they do not make contact with the DNA backbone, their role in site-specific recognition could be similar. To our knowledge, no region analogous to the one we discovered in VqmA_Phage_ has been shown to confer promoter specificity to a transcription factor. A caveat of our interpretation is that, as noted, we do not have a structure of VqmA_Phage_ and we mapped the residues identified in our VqmA_Phage_ mutagenesis to the DPO-VqmA_Vc_-DNA crystal structure. Therefore, it remains possible that the residues we identified here do indeed make contacts with DNA. Going forward, determining the structure of VqmA_Phage_ bound to DNA should reveal the mechanism that confers recognition of P*qtip* and the role that these residues play, individually and collectively, in determining DNA-binding specificity.

Genomic sequencing data have revealed the presence of many QS receptor-transcription factors encoded in phage genomes (24). In general, however, their transcriptional outputs are uncharacterized, with the exception of VqmA_Phage_, which is promiscuous with respect to binding to P*vqmR* and P*qtip*, the only two promoters tested to our knowledge. It remains possible that VqmA_Phage_ regulates additional genes specifying bacterial and or/phage functions. Given that VqmA_Phage_ can regulate biofilm formation through its control of *V. cholerae vqmR*, probing the host regulon controlled by VqmA_Phage_ under various growth conditions could reveal unanticipated roles of QS in phage-*Vibrio* interactions.

Finally, we found that the VqmA_Vc_^A192E^ variant exhibited modest, but detectable binding to P*qtip*, whereas the VqmA_Vc_ quadruple mutant, and the VqmA_Vc_ ^*171-199^ mosaic protein did not. Western blot and P*vqmR-lux* assays eliminated the possibility that any of the mutant proteins were not expressed or were misfolded. Rather, we infer that a particular regional conformation in the VqmA proteins is required for this key residue to function properly. Our results also show that exchanging both the VqmA_Vc_ HTH motif and C-terminal 25 residues with the corresponding residues from VqmA_Phage_ enables some but not WT-level binding to P*qtip*. This finding supports the notion that a set of non-contiguous amino acids or a particular conformation of the VqmA_Vc_ DBD prevents binding to P*qtip*. This arrangement is perhaps not surprising given that *V. cholerae* would pay a significant penalty if VqmA_Vc_ bound the phage VP882 *qtip* promoter, as the consequence would be the launch of the phage lytic program and death of the host cell. To our knowledge, VqmA_Vc_ binds to only one promoter, P*vqmR* (3). Thus, even in the context of the *V. cholerae* genome, VqmA_Vc_ transcriptional activity is tightly constrained. It is possible that other negative ramifications stem from non-specific VqmA_Vc_ binding in the *V. cholerae* genome. Distinct mechanisms are employed to restrict other QS receptor/transcription factors from promiscuously binding to DNA. For example, LuxR-type QS receptors can typically bind >100 promoters, but their solubilization, stability, and DNA-binding capabilities strictly rely on being bound to an AI whose availability is, in turn, highly regulated (25–29). Therefore, precise control of gene expression is maintained in many QS circuits by confining QS receptor activity to the ligand-bound form coupled with discrete affinities of the ligand-receptor complexes for target promoters. By contrast, VqmA_Vc_ is expressed constitutively, and its DNA-binding capabilities are not limited by the presence of an AI. Thus, exquisitely tight control over promoter DNA-binding specificity by VqmA_Vc_ --restricting it to one and only one promoter--is apparently crucial for proper regulation of gene expression and survival.

## MATERIALS AND METHODS

### Bacterial strains, plasmids, primers, and reagents

Strains, plasmids, primers, and gBlocks used in this study are listed in Tables 1-4, respectively. In all experiments, Δ*tdh V. cholerae* and Δ*tdh E. coli* strains were used except in the experiment assaying expression of P*vqmR-lux* and P*qtip-lux* in response to the DBD_Vc_, DBD_Phage_, GST-DBD_Vc,_ and GST-DBD_Phage_ proteins. In that case, the *E. coli* strain contained the WT *tdh* gene. *V. cholerae* and *E. coli* were grown aerobically in lysogeny broth (LB) at 37°C. Antibiotics and inducers were used at the following concentrations: 50 units mL^-1^ polymyxin B, 200 µg mL^-1^ ampicillin, 5 µg mL^-1^ chloramphenicol, 100 µg mL^-1^ kanamycin, 0.2% arabinose, and 1 mM Isopropyl β-D-1-thiogalactopyranoside (IPTG).

Primers were obtained from Integrated DNA Technologies. Gibson assembly, intramolecular reclosure, and traditional cloning methods were employed for all cloning. PCR with Q5 High Fidelity Polymerase (NEB) was used to generate insert and backbone DNA. Gibson assembly relied on HiFi DNA assembly mix (NEB). All enzymes used in cloning were obtained from NEB. Mutageneses of the VqmA_Phage_ and VqmA_Vc_ DBDs was accomplished using the GeneMorph II EZClone Domain Mutagenesis Kit (Agilent) according to the manufacturer’s instructions. Transfer of plasmids carrying *vqmA* genes into the *V. cholerae* phage VP882 lysogen employed conjugation followed by selective plating on polymyxin B, chloramphenicol, and kanamycin, based on previously described protocols (30).

### Genetic screens for VqmA_Phage_ and VqmA_Vc_ DNA-binding mutants

*E. coli* carrying a library of plasmid-borne *vqmA*_*Phage*_ mutants was mated with *V. cholerae* harboring a phage VP882 mutant (*vqmA*_*Phage*_::Tn*5*) and the P*vqmR-lux* reporter integrated at the *lacZ* locus. Exconjugant *V. cholerae* colonies were collected and streaked onto LB agar plates supplemented with polymyxin B, chloramphenicol, kanamycin, and arabinose. P*vqmR-lux* activity of surviving exconjugants was assayed using an ImageQuant LAS4000 imager (GE). *V. cholerae* colonies that produced light were harvested for plasmid DNA preparation. Isolated plasmid DNA was subsequently transformed into *E. coli* strains carrying P*qtip-lux* or P*vqmR-lux* to validate activity.

A library of plasmid-borne *vqmA*_*Vc*_ mutants was transformed into *E. coli* carrying the P*qtip-lux* reporter. Transformants were plated on LB agar supplemented with ampicillin, kanamycin, and arabinose. P*qtip-lux* activity was assayed using an ImageQuant LAS4000 imager.

### Growth, lysis, and bioluminescence assays

To measure growth of *V. cholerae* phage VP882 lysogens or activation of the P*vqmR-lux* and P*qtip-lux* reporters in bacterial strains, overnight cultures of *V. cholerae* or *E. coli* were back-diluted 1:1000 into LB medium supplemented with appropriate antibiotics prior to being dispensed (200 µL) into 96-well plates (Corning Costar 3904). Arabinose was added as specified. The plates were shaken at 37°C and a Biotek Synergy Neo2 Multi-Mode reader was used to measure OD_600_ and bioluminescence. For bioluminescence assays, relative light units (RLU) were calculated by dividing bioluminescence by the OD_600_ after 5 h.

### Protein expression, purification, and electrophoretic mobility shift assay (EMSA)

Protein expression and purification were performed as described (9). EMSAs were performed as described (8) with the following modifications: following electrophoresis, 6% DNA retardation gels were stained with SYBR Green (Thermo) and visualized using an ImageQuant LAS 4000 imager with the SYBR Green settings. Unless specified otherwise, the highest concentration of VqmA assessed was 600 nM. 25 nM P*vqmR* or P*qtip* DNA was used in all EMSAs. The percentage of promoter DNA bound was calculated using the gel analyzer tool in ImageJ and the estimated EC_50_ values were derived from EC_50_ analyses in Prism.

### Western blot analysis

Western blot analyses probing for abundances of 3xFLAG-tagged proteins were performed as reported (3) with the following modifications: *E. coli* and *V. cholerae* carrying N-terminal 3xFLAG-tagged VqmA_Vc_ and N-terminal 3xFLAG-tagged VqmA_Phage_ alleles were back-diluted 1:1000 in LB supplemented with appropriate antibiotics and harvested after 6 h and 4 h of growth at 37°C, respectively. Cells were resuspended in Laemmli sample buffer at a final concentration of 0.006 OD/µL. Following denaturation for 15 min at 95°C, 5 µL of each sample was subjected to SDS-PAGE gel electrophoresis. RpoA was used as the loading control (Biolegend Inc.). Signals were visualized using an ImageQuant LAS 4000 imager.

### Sequence alignments

Protein and DNA sequences in FASTA format were aligned in the BioEdit Sequence Alignment Editor using the default setting under the ClustalW mode. Figure 2A and Supplementary Figure 1 were prepared via the ESPript 3.0 online server (31).

## ACKNOWLEDGEMENTS

We thank members of the Bassler laboratory for insightful discussions. This work was supported by the Howard Hughes Medical Institute, National Institutes of Health Grant R37GM065859, and National Science Foundation Grant MCB-1713731 (BLB), NIGMS T32GM007388 (OPD), a Charlotte Elizabeth Procter Fellowship provided by Princeton University, and a National Defense Science and Engineering Graduate Fellowship supported by the Department of Defense (JES). The content is solely the responsibility of the authors and does not necessarily represent the official views of the National Institutes of Health.

